# Tapeworm infection affects sleep behavior in three-spined stickleback

**DOI:** 10.1101/2024.01.30.577962

**Authors:** Marc Bauhus, Sina Mews, Joachim Kurtz, Alexander Brinker, Robert Peuß, Jaime Anaya-Rojas

## Abstract

Sleep is a highly complex and conserved biological process that affects several body functions and behaviors. Recent evidence suggests that there is a reciprocal interaction between sleep and immunity. For instance, fragmented sleep can increase the probability of parasitic infections and reduce the ability of infected hosts to fight infections. It is also known, particularly in humans and other mammals, that viral and bacterial infections alter the sleep patterns of infected individuals. However, the effects of macro-parasitic infections on sleep remain largely unknown. In this study, we investigated whether macro-parasite infections could alter the sleep of their hosts. We experimentally infected three-spined sticklebacks (*Gasterosteus aculeatus*) with the tapeworm *Schistocephalus solidus* and used a hidden Markov model to characterize sleep-associated behaviors in the sticklebacks. At an early time-point, 1-4 days after parasite exposure, infected fish showed no difference in sleep compared with non-exposed fish, whereas fish that were exposed but could fend off the infection slept less during the daytime. At a later time-point, 29-32 days after exposure, infected fish slept more than uninfected fish, while exposed-but-not-infected fish slept less than non-exposed fish. Using RNA-seq of brain tissue, we identified several immune- and sleep-associated genes that potentially underlie the observed behavioral changes. These results provide the first insight into the complex association between macro-parasite infection, immunity, and sleep in fish and may thus contribute to a better understanding of the reciprocal interaction between sleep and immunity.

## Results and Discussion

Sleep is a fundamental biological process in animals (Campbell and Tobler, 1984) and is characterized by periods of behavioral inactivity that are easily reversible (Tononi, 2000; Yokogawa et al., 2007). Current hypotheses assume that sleep has evolved to maintain brain connectivity, plasticity, and health, and to reduce caloric consumption (Krueger et al., 2016; Lesku and Schmidt, 2022), and is regulated by neurological processes (Brown et al., 2012), circadian rhythms, and duration of wakefulness (Borbély et al., 2016). Furthermore, parasitic infections can affect the regulation of neuropeptides such as hypocretin, which can promote wakefulness and influence the cellular composition of the immune system during myelopoiesis (McAlpine et al., 2019). Although the impact of bacterial and viral infections on sleep-immunity interactions has been extensively studied in mammals (Fang et al., 2016; Kimura-Takeuchi et al., 1992; Toth, 2019; Toth and Kreuger, 1988), the influence of macro-parasite infections on such interactions remains largely unexplored.

In this study, we aimed to address this gap by examining the effects of a macro-parasite infection on sleep duration in a fish hosts. For this, we made use of an increasingly studied natural host-parasite system that allows for experimental infection in the laboratory, the three-spined stickleback (*Gasterosteus aculeatus*) and its host-specific tapeworm *Schistocephalus solidus* (Barber and Scharsack, 2010; Demandt et al., 2020; Hammerschmidt and Kurtz, 2009; Scharsack et al., 2021; Weber et al., 2022) (Figure 1A). We experimentally infected sticklebacks and measured the effects of the infection on the overall activity of individuals during two 4-day-periods, 1-4 days and 29-32 days after parasite exposure (dpe) (Figure 1B, C). In sticklebacks, *S. solidus* grows strongly and manipulates the host’s behavior to increase the chances of transmission to the final host (Barber et al., 2004; Talarico et al., 2017). Due to an increasing understanding of *S. solidus* biology and its effects on stickleback behavior and immunity, this host-parasite system is well-suited to study the effects of macro-parasitic infections on sleep behavior in vertebrate hosts.

**Figure 1.**
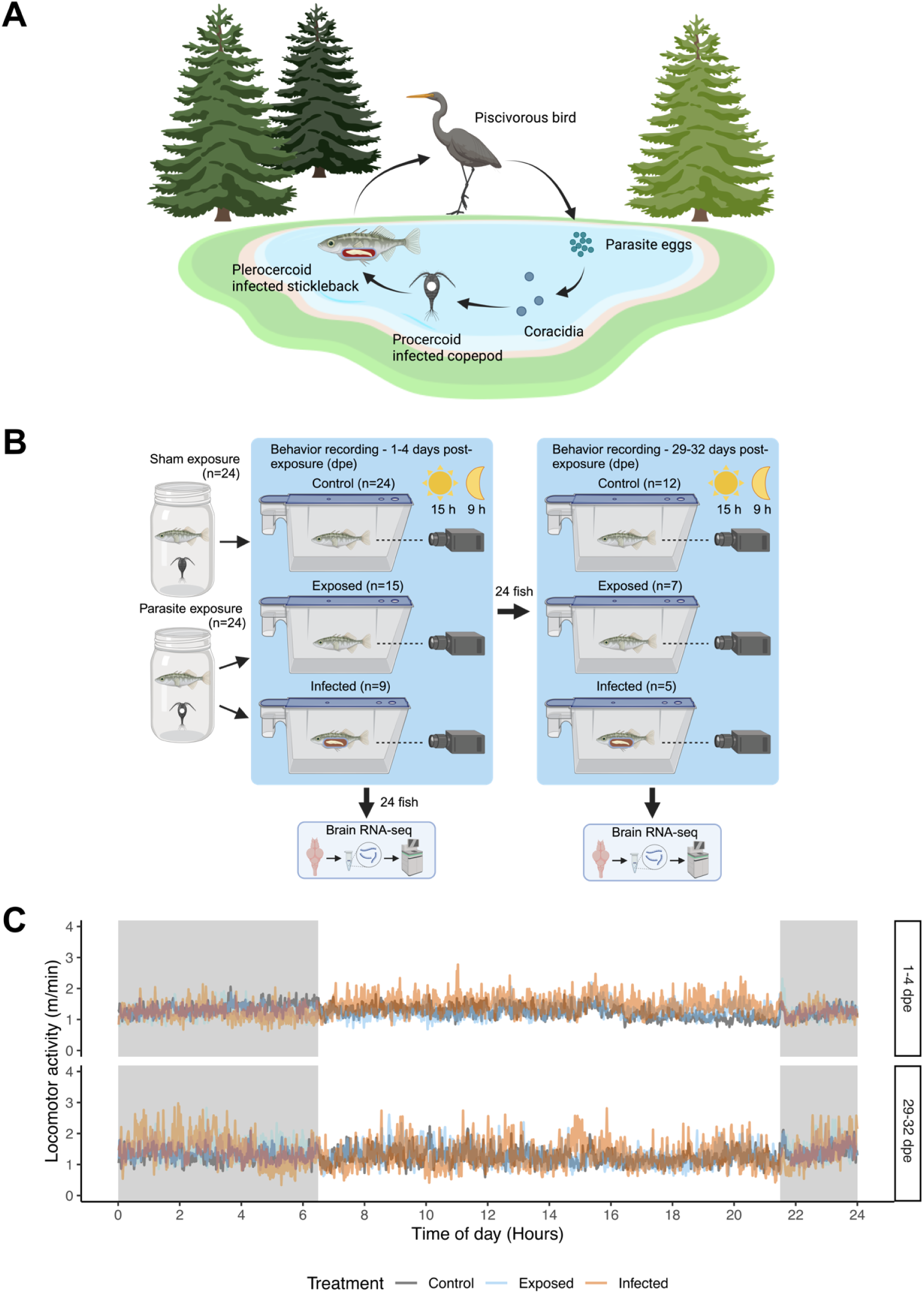
Tracking locomotor activity of three-spined stickleback to estimate sleep upon infection with *S. solidus*. (A) Lifecycle of *S. solidus*, a trophically transmitted tapeworm with two intermediate hosts – cyclopoid copepods and three-spined sticklebacks – and warm-blooded fish-eating animals (mostly birds) as final hosts, where the worms reproduce (Barber and Scharsack, 2010). (B) Experimental design. Individual sticklebacks were exposed to either parasite-naive copepods or *S. solidus*-infected copepods (indicated by white dots). The resulting non-exposed (control), exposed but not infected (exposed), and infected (infected) fish were video-recorded 1-4 and 29-32 days post parasite exposure (dpe). Half of the fish were dissected after the first recording, whereas the other half were recorded at both time points and then dissected. All fish were inspected for their infection status, and brain tissue was used for RNA-seq, providing transcriptomic data for the time points 4 and 32 dpe. (C) Mean locomotor activity (m/min) of the control (black), exposed (blue), and infected (red) fish averaged over 24 h for both periods (1-4 and 29-32 dpe). The figure was created using Biorender.com.

Using hidden Markov models (HMMs), we identified three distinct behavioral states in our experimental fish: a sleep-like behavioral state (sleep), a state of moderate activity, and a state of high activity (Figure 2A). We compared the proportion and probability of sleep between three types of experimental fish: control (fish that were not exposed to *S. solidus*), exposed (fish that were exposed but not infected), and infected (Figure 1B). We predicted that during the early stage of infection (1-4 dpe), both infected and exposed fish would sleep less than control fish due to the immune response against the parasite (Drake et al., 2000; Toth and Kreuger, 1988). We further predicted that at a later stage of infection (29-32 dpe), infected fish might sleep more because of the energy demand of the parasite, thereby increasing the need for more efficient energy conservation (Schmidt, 2014). We sequenced the transcriptome of fish brains at four and 32 dpe to identify potential mediators of the sleep-immunity interactions triggered by *S. solidus* infection.

**Figure 2.**
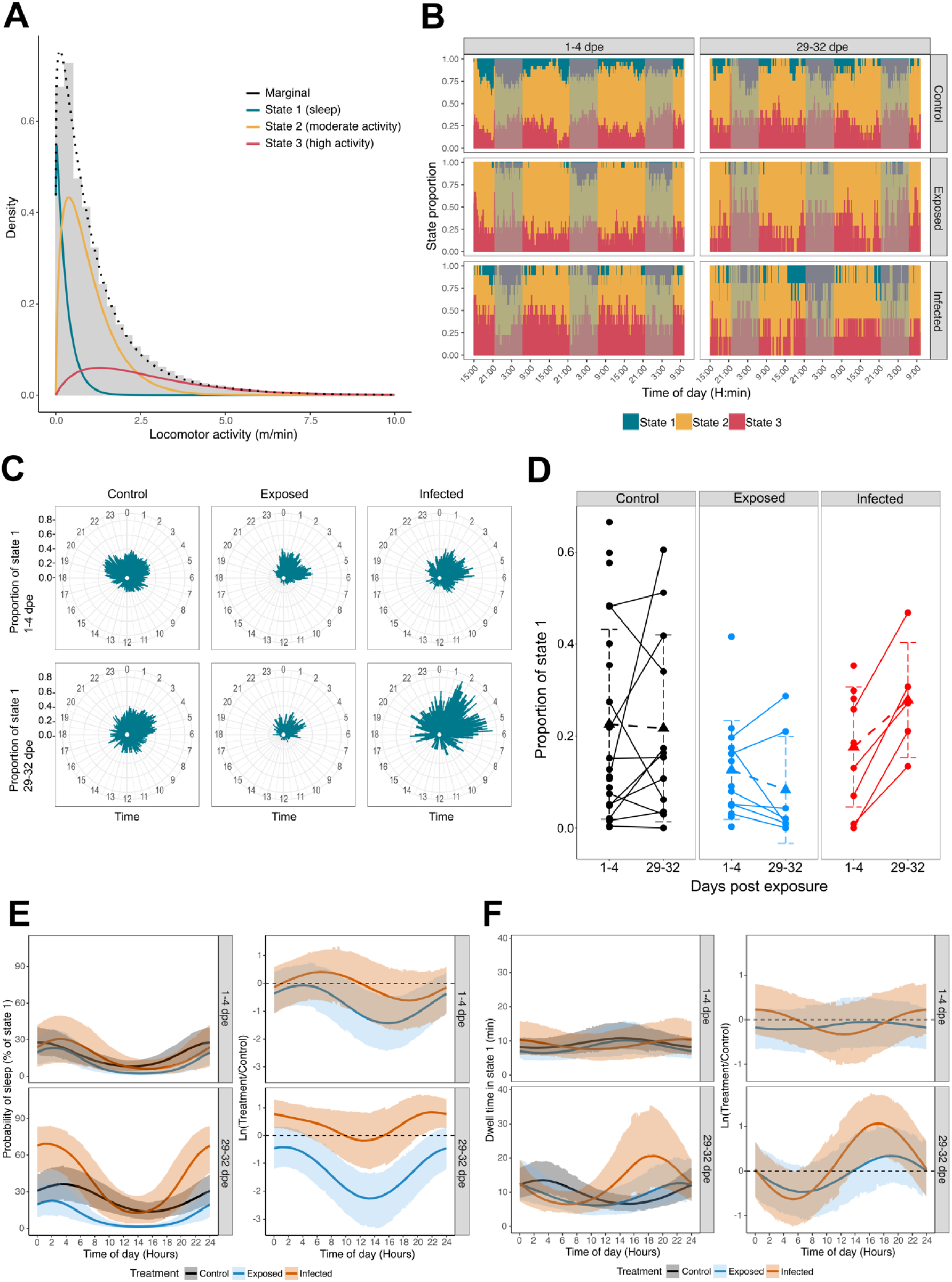
Stickleback sleep increased upon *S. solidus* infection 29-32 days post parasite exposure. (A) Histogram of three different behavioral states (state 1=sleep, state 2=moderate activity, and state 3=high activity) identified by the hidden Markov model fitted to the observed locomotor activity. The state-dependent distributions were weighted according to the proportion of time spent in different states, and the dashed line indicates the associated marginal distribution under the fitted model. (B) Activity patterns of control, exposed, and infected fish 1-4 and 29-32 days post-exposure (dpe). The proportions of time spent in each state are shown over the respective recording time span. Nighttime is indicated by shaded areas. (C) Proportions of time spent in state 1 in control, exposed, and infected fish averaged over 24 h per recording time (1-4 and 29-32 dpe). (D) Individual proportion of time spent in state 1 1-4 and 29-32 dpe. Data points of individuals that were recorded repeatedly were connected by a line to show within-individual development of sleep over time. The triangles indicate mean values. The dashed lines connecting the means between 1-4 and 29-32 dpe show the mean development of sleep over time in the respective treatments. The dashed error bars represent the standard deviation. (E) On the left side: Probabilities (i.e., expected percentages) for control, exposed, and infected fish occupying state 1, corresponding to the periodic stationary distribution of the HMM per recording time (1-4- and 29-32 dpe). The middle lines display the mean probabilities, and the upper and lower areas represent 95% CIs. On the right side, the logarithmic ratio of the deviation of exposed and infected fish from the respective control (dashed line) was derived from simulations based on the HMM. The middle lines display the means and the upper and lower areas with 95% confidence intervals of the distribution. (F) On the left side: Expected dwell times (i.e., time spent continuously in one state) as a function of time of day for control, exposed, and infected fish in state 1 per recording time. The middle lines display the mean dwell times and the upper and lower lines represent the respective 95% confidence intervals. On the right side: Logarithmic ratio of the deviation of exposed and infected fish from the respective control (dashed line) derived from simulations based on the HMM. The middle lines display the means and the upper and lower areas of the 95% CIs of the distribution.

Overall, we found that sticklebacks spent approximately 19% of their time sleeping, mostly during the night (i.e., between 21:30 and 6:30, Figure 2B and C). We found partial support for our predictions. Contrary to our first prediction, we observed only weak differences in sleep between 21:30 and 6:30 among the three types of fish early after exposure (1-4 dpe) (Figure 2B–E). Specifically, the probability of finding infected fish sleeping during this period was approximately 1.3% higher than that of the control fish (mean = 1.3%; 95% CI: −20.9 to 27.6% of sleep) and 7% higher compared to that of exposed fish (mean = 7%; 95%; CI: −14.0 to 31.8%; Figure 2E).

These results suggest little impact of the tapeworm infection on sleep, perhaps because *S. solidus* can effectively evade the immune response of sticklebacks (Hammerschmidt and Kurtz, 2005; Scharsack et al., 2007). This hypothesis is supported by our brain transcriptome analysis, which shows that there was only one immune response-related GO-term among the most significantly enriched GO-terms (‘positive regulation of cytokine production,’ Figure 3D). However, some immune-related genes were differentially expressed. For example, colony stimulating factor 1 receptor a (CSF1Ra), which is involved in neuroinflammation and monocyte (microglia) differentiation and proliferation (Chitu et al., 2016), was upregulated in infected fish at 4 dpe (Table 1).

**Figure 3.**
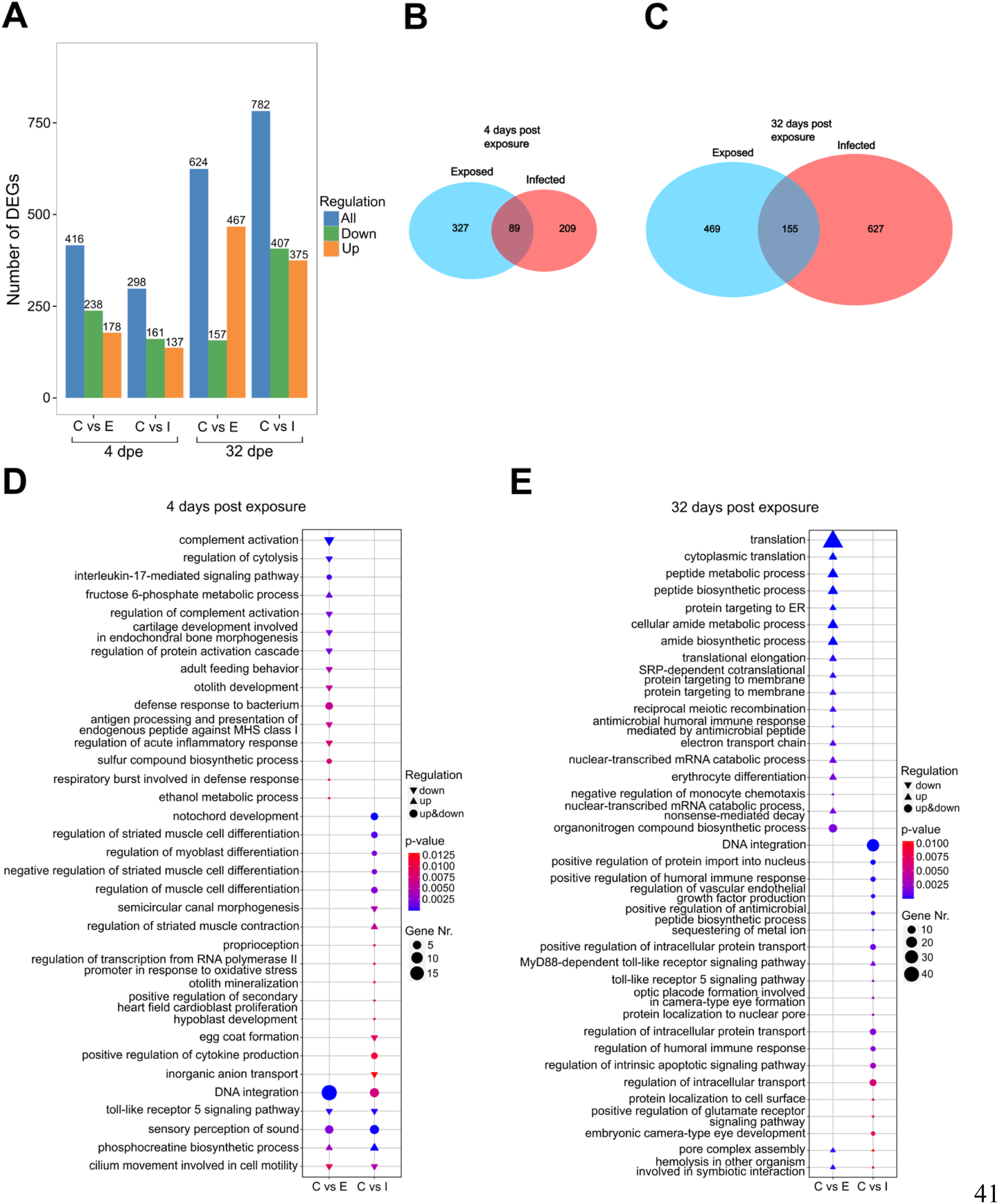
Differential gene expression patterns and GO-terms between Control, Exposed, and Infected fish. (A) Number of differentially expressed genes (DEGs) (total, up-, and downregulated) of the control (C) versus exposed (E) and infected (I) fish at 4 and 32 dpe. (B) Venn diagram showing the number of unique and shared DEGs between Exposed and Infected fish versus control fish at 4 dpe (C) and 32 dpe. (D) The 20 most significantly enriched and non-redundant Gene Ontology (GO) terms between control and exposed but not infected and infected fish at 4 dpe (E) and 32 dpe. The shape indicates the direction of the regulation. The size of the shape shows the number of DEGs expressed in this GO term and the color gradient represents the significance of enrichment

**Table 1.**
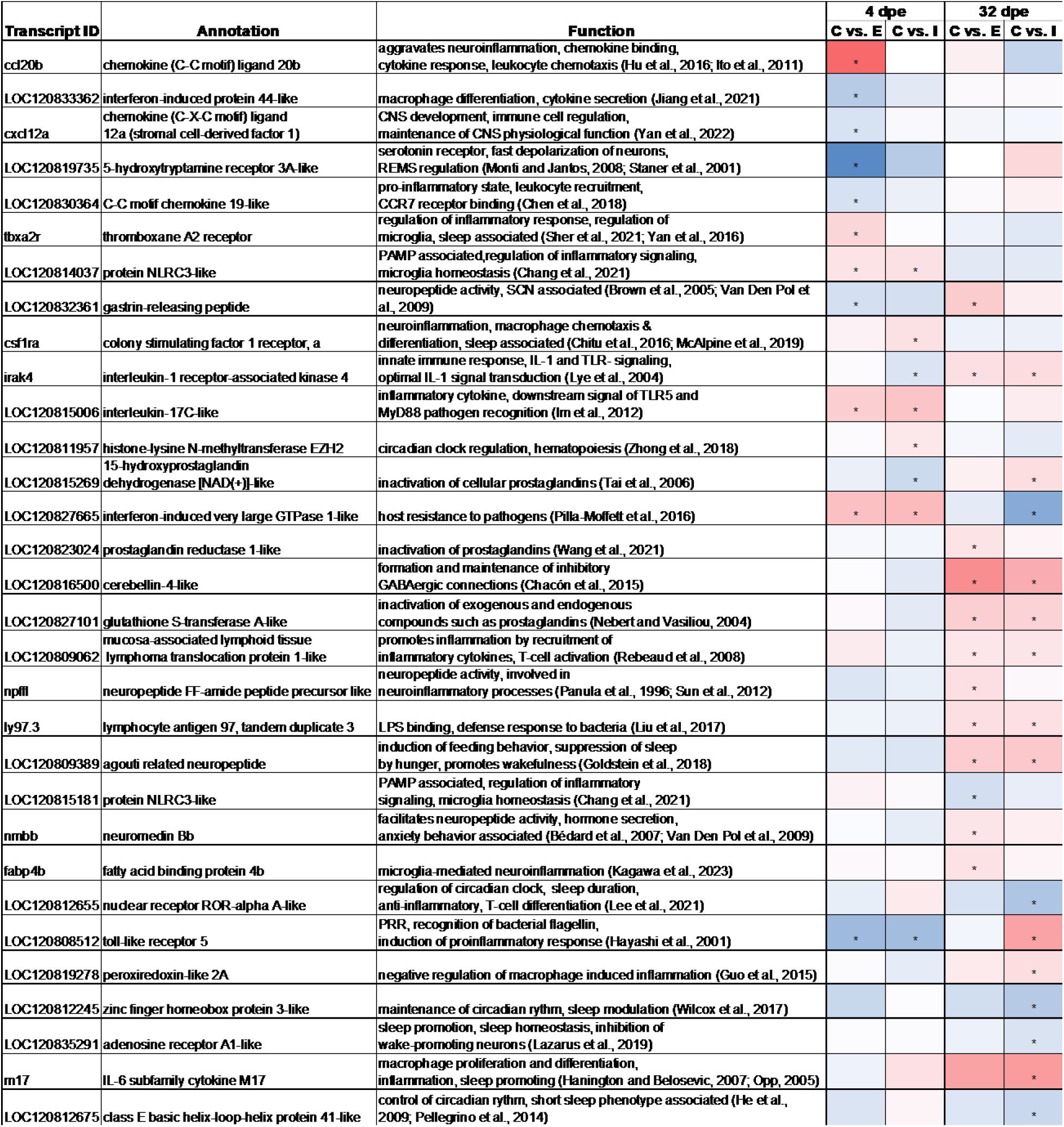
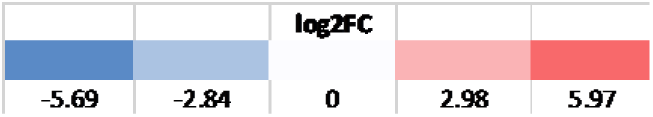
Immune- and sleep-associated genes are differentially expressed in exposed and infected fish. Table containing the NCBI transcript ID, annotation, short summary of (putative) biological function, and heat map showing the log2 fold change of each differentially expressed gene per treatment (Control=C, Exposed=E, Infected=I) and time point (4 and 32 dpe) Significant up- or down-regulation is indicated with an asterisk (log2FC>1, p</=0.05)

Interestingly, our modeling approach also suggests that sticklebacks can be found sleeping during the day and that parasite exposure may impact this behavioral pattern. For instance, we found that exposed fish had an approximately 7% lower sleep probability during the daytime (6:30-21:30) than control fish at 1-4 dpe (95%CI: −18.6 to 2.6%; Figure 2E). These differences could be the result of immune responses to the parasites. The brain transcriptomic analysis revealed that the chemokine (C-C motif) ligand 20b (ccl20b) was upregulated in exposed fish compared to that in control fish (Table 1). As ccl20b is involved in neuroinflammation, chemokine binding, cytokine response, and leukocyte chemotaxis (Hu et al., 2016; Ito et al., 2011), the observed upregulation of this gene may indicate an immune response against the parasite or translocated gut bacteria following gut perforation by the parasite (Hahn et al., 2022). However, whether these or other genes involved in neuroinflammation are involved in the observed changes in sleep probability of exposed fish needs to be functionally validated.

As predicted for the later stage of infection (29-32 dpe), we found that infected fish slept more than control fish (Figure 2B, 2C), especially during the night hours (21:30-06:30). Overall, the probability of sleep during the night for infected fish increased from approximately 26.4% (95%CI: 7.3 to 46.7%) to 62.5% (95%CI: 39.1 to 82.2%); during the day, it increased from 12% (95%CI: 1.5 to 28.9%) to 26.0% (95%CI: 4.8 to 56.1%; Figure 2E). Specifically, during the night, the probability of sleep for infected fish 29-32 dpe was approximately 30% higher than that of control fish (95%CI: 3.6–54.9% of sleep) and 43% higher than that of exposed fish (95%CI: 17.6–65.9% of sleep). Furthermore, the time that the infected fish spent sleeping uninterruptedly during the day (i.e., the expected time spent in state 1 before switching to a different state) was longer than that at night. During the nights, infected fish spent approximately 10 min sleeping uninterruptedly (95%CI: 5.7 to 15.4 mins) 1-4 dpe and 11.5 mins (95%CI: 4.3 to 21.4 mins) 29-32 dpe. However, during the day, the differences in the time spent sleeping uninterruptedly changed from approximately 8.9 mins (95%CI: 4.5 to 14.1 min) to 14.07 mins (95%CI: 3.9 to 21.1 min) from 1-4 dpe to 29-32 dpe (Figure 2F). Interestingly, during the second part of the day (i.e., from 12:00 to 21:00) infected fish spent almost 10 min more sleeping uninterruptedly than control fish (95%CI: −0.8 to 24.3 mins) and approximately 8.8 mins more than exposed fish (95%CI: −3.0 to 22.7 mins; Figure 2F).

Within the first 12 weeks of infection, *S. solidus* shows a steep increase in weight gain (Barber and Svensson, 2003). If the energy cost of *S. solidus* is high, infected fish require more sleep to compensate for their energy requirements. There is ample evidence to support this hypothesis. For example, *S. solidus* drains a significant amount of energy from its stickleback host, particularly during single infections (Nordeide and Matos, 2016). Additionally, when infected with large intestinal parasites, sticklebacks often specialize in eating many smaller prey items or selectively preying on a few larger items to compensate for the energetic cost of infection (Molbert and Goutte, 2022). Hence, infected sticklebacks could cope with the cost of infection by sleeping more (Schmidt, 2014).

The increased sleep of infected fish at 29-32 dpe could also be partially explained by the direct and indirect effects of the immune evasion strategies of *S. solidus*. Immune evasion is a key strategy used by pathogens to improve their fitness (Schmid-Hempel, 2008). For trophically transmitted parasites, such as *S. solidus*, the energy invested in immune evasion has to trade-off with somatic growth; therefore, the increased sleep observed in the later infection stage could be a way to modulate the host immune system to enhance nutrient gain for the parasite (Schmid-Hempel, 2008).

Immune responses against bacterial or viral infections, as well as the experimental inoculation of inflammatory cytokines, can have sleep-promoting effects, depending on the infection dose and stage (Besedovsky et al., 2019; Fang et al., 2016; Kubota et al., 2001b, 2001a; Mullington et al., 2000; Opp, 2005; Toth and Kreuger, 1988). This is thought to contribute to a more efficient defense against parasites (Preston et al., 2009; Imeri and Opp, 2009). Therefore, this might also be true for long-term infections with macroparasites such as tapeworms. Although successful clearance of the tapeworm is unlikely to occur late during infection, immunity may reduce parasite growth (Scharsack et al. 2007). We found that both infected and exposed fish had more differentially expressed genes (DEGs) than the control fish at the late (32 dpe) compared to the early (4 dpe) time point (Figure 3A). Among these DEGs, we identified several genes with possible associations with immune response and sleep regulation (Table 1). For example, the anti-inflammatory and circadian rhythm-associated nuclear receptor ROR-alpha A-like (RORA A-like) (Lee et al., 2021) was downregulated in infected fish, whereas several inflammation-associated genes were upregulated (Table 1), which may have contributed to the observed increase in sleep during night hours.

*S. solidus*, such as many other trophically transmitted parasites with complex life cycles, is known to manipulate the anti-predatory behavior of its stickleback host to increase the likelihood of transmission to the final host (Barber et al., 2004). Therefore, the parasite might make use of immune-sleep interactions in its host by utilizing the increased arousal threshold during sleep to increase the predation risk of the stickleback (Barber et al., 2004; Krueger et al., 2016). In fact, we observed that infected fish spent, on average, more time continuously in a sleep state (dwell time) during the daytime (Figure 2F), when the predation risk by birds is thought to be higher (Quinn et al., 2012; Stumbo and Poulin, 2016). In a recent brain transcriptomics study using stickleback after three month of infection with *S.solidus*, it was suggested that the inositol pathway which is involved in human neuropsychiatric diseases (Frej et al., 2017) might be involved in the behavioral manipulation by the parasite (Grecias et al., 2020). But we did not find a significant differential expression of the associated gene inositol monophosphatase 1 (impa1) at 1-4 or 29-32 dpe. However, it is likely that the parasites in our experiment were not yet infective to the final host, because they were below the published threshold mass for reproduction in the final host of ∼50 mg (Barber et al., 2004; Supplementary Table 1). Nevertheless, it is possible that this threshold is lower for some parasite populations. Therefore, further experiments at later stages of infection are needed to test whether the fish show more sleep than in our experiment.

We also found that exposed fish showed an overall lower sleep proportion (Figure 2B, 2C) and a lower probability of sleep during the daytime (Figure 2E) compared to control fish 29-32 dpe. Interestingly, these fish also showed a higher number of upregulated GO terms and DEGs associated with translation and protein transport (Figure 3E). We also identified upregulated genes that were directly related to neuroinflammation, neuropeptide activity, and sleep regulation (Table 1). For example, interleukin-1 receptor-associated kinase 4 (irak4) facilitates the inflammatory response (Lye et al., 2004), and agouti-related neuropeptide promotes wakefulness in mice (Goldstein et al., 2018). The latter could be a candidate gene responsible for the reduction in sleep behavior of exposed fish 29-32 dpe, although this gene was also upregulated in infected fish. However, the few shared DEGs between exposed and infected fish could indicate that immunological and associated (neuro-) inflammatory processes differed substantially between these groups (Figure 3B, 3C). Differences between exposed and infected fish have also been found in other studies, even at later stages post-exposure, such as changes in liver gene expression, body condition, and gut microbiome composition (Hahn et al., 2022; Piecyk et al., 2019; Scharsack et al., 2021), which might indicate chronic inflammation following an immune response to either the parasite or translocated bacteria in exposed fish.

However, whether immune responses play a role in the observed effects of *S. solidus* infection on sleep and the genetic and cellular pathways involved need to be further investigated. Combining analyses of the immune response at the hematopoietic site of the fish, the head kidney, with brain RNA-seq could reveal more detailed insights into the hypothesized immune-sleep interaction. Furthermore, it is necessary to gain a better understanding of the sleep ecology of three-spined stickleback. To date, only two studies have explored rhythmicity in activity behavior during the breeding season (Sevenster et al., 1995) and overall circadian rhythmicity (Brochu and Aubin-Horth, 2021). In contrast to our findings, the latter study found stickleback to be nocturnal. However, they also observed striking individual differences in activity behavior, which is in line with our findings.

The differences in diurnal activity and sleep among individuals might be due to the large variety of ecological conditions to which sticklebacks are adapted in nature, which has also been discussed for other animals (Rattenborg et al., 2017). Therefore, sleep may also be an important phenotypic mechanism for individual, temporal niche conformance under fluctuating environmental conditions (Trappes et al., 2022). Nevertheless, our study is the first to specifically focus on sleep-like behavior in this system. Because of the limitations of electroencephalographic slow-wave measurements in fish, which is a common method for characterizing sleep in mammals and birds (Patel et al., 2021; Rattenborg et al., 2017), our activity-based characterization of sleep with a hidden Markov model provides an objective approach for sleep research, even in non-model organisms. This could be extended by measuring the arousal threshold of inactive individuals (Yokogawa et al., 2007; Yoshizawa et al., 2015) to gain more certainty regarding the observed behavioral state. Therefore, investigating our observed infection-induced changes in sleep-like behavior together with more detailed information about the sleep of fish in general could help to better understand the possible ecological and evolutionary consequences of the hypothesized immune-sleep interaction.

## Conclusion

Our study showed that macro-parasite infections can affect the sleep of the infected fish and that these effects become stronger as the infection progresses. Moreover, we identified differentially expressed genes in the brains of the fish that are associated with immune responses and sleep regulation, which might be involved in the observed changes in sleep. These changes might have interesting ecological and evolutionary implications, which can be further explored by future studies focusing on later time points post-exposure, as well as on more detailed insights into immune response dynamics and sleep of animals to deepen our understanding of the hypothesized immune-sleep interaction.

## Ethics

All animal experimental procedures were approved by and executed in accordance with the local veterinary and animal welfare authorities (project number: 84-02.04.2014.A368)

## Acknowledgements

We thank Luis Garcia Rodriguez, David Martín Fernández, and Jacomo Krause from the workshop of the Faculty of Biology, University of Münster, for their support in building and programming the automated behavior recording system. We also thank Andreas Revermann for helping with fish collection. We also acknowledge the help of Michelle Borgers and Britta Bock in fish dissection and parasite screening. Furthermore, we thank Britta Bock and Kathrin Brüggemann for their assistance with RNA-extractions. We also acknowledge Ilka Rauch for her support with copepod infections, and Georg Plenge for technical support with the automated behavior recording system. We appreciate the help of Carlina C. Feldmann in providing helpful advice on HMM model fitting and plotting. This work was supported by institutional funding to J.K. and funded by the German Research Foundation (DFG) as part of the CRC TRR 212 (NC³), projects C05 (granted to J.A.R. and J.K., Project number 471674348), and D06 (granted to Roland Langrock, Project number 396782756).

## Author contributions

MB, J.K, J.A.R., and R.P. conceived the project. M. B. performed the infections and sleep experiments with support from J.A.R., R.P., J.K., and A.B.. S.M. performed the HMM analysis with support from J.A.R.. M.B. performed the transcriptomic analysis with support from R.P.. M.B. wrote the first draft of the manuscript, and all authors read, contributed to, and approved the final version of the manuscript.

## Declaration of interests

The authors declare that they have no conflicts of interest.

## Data and code availability

The data and code will be made public upon the acceptance of the manuscript by a public server.

## Material & Methods

### Experimental stickleback population and husbandry

For this experiment, we used F1 offspring of wild-caught three-spined sticklebacks from Lake Constance which are commonly infected with *S. solidus* (Baer et al., 2022). Parental fish were caught in late May 2021 using minnow traps in a river mouth and marina close to Langenargen, Germany, with permission from the Fisheries Research Station Langenargen. After 2-3 weeks of acclimatization to laboratory conditions, we collected ovaries and sperm from the wild-caught fish and artificially fertilized them. The offspring were raised in tap water at 17°C and fed daily with freshly hatched *Artemia* larvae and frozen lobster eggs. After reaching a size of approximately 20 mm, we transferred the fish from smaller breeding tanks to 14 liter aquaria with recirculating tap water, temperature control, mechanical, UV, and biological filtration (Vewa Tech, Hamm, Germany). Fish were fed frozen lobster eggs and chironomid larvae. At 4 months of age, the fish were fed only chironomid larvae. Over their entire lifespan, sticklebacks were exposed to a 15/9 h light/dark cycle with 1h of simulated sunset and sunrise. Fish were maintained in their respective families (siblings derived from one mating pair).

### Infection with *S. solidus*

We experimentally infected sticklebacks with *S. solidus* from a stream stickleback population residing in the Ibbenbürener Aa, northwest Germany, to remove any potential effects of local adaptation. We used eggs from one breeding pair (IBB 26) for infections, which were artificially bred according to Schärer and Wedekind (1999). The eggs were incubated in Petri dishes for at least two weeks at 20°C in the dark. We isolated male copepods from a laboratory stock of *Macrocyclops albidus* in 24-well plates two days before parasite exposure. We starved the copepods during isolation to increase the probability of parasite consumption. We induced hatching of the parasites by 3 h of light exposure in the evening, followed by 9 h of darkness and subsequent light exposure in the morning. Two to three free-swimming coracidia were collected using a pipette and transferred to each isolated copepod. Per behavioral recording of sticklebacks, we exposed 36 copepods to *S. solidus* and eight remained unexposed in the control group. After exposure, the copepods were incubated at 20°C and fed every 48 h with 10-20 live paramecium per copepod. The copepods were maintained for at least 12 d to allow the parasite to develop into the infective procercoid stage (Benesh and Hafer, 2012). We then screened the copepods for procercoids under a microscope and used single-infected copepods for stickleback infection. Five families of F1 Lake Constance sticklebacks were used for the experiment. We exposed one fish per family and recorded their behavior, and the other remained unexposed for the control. At the time of exposure, the fish were between 4,5 and 6,5 months old. For each behavioral recording, we isolated 8-10 fish in jars with 400 ml block water. After 24 h of acclimatization, we exposed each 4-5 sticklebacks to an infected or non-exposed copepod in the jar (Figure 1). Two days before parasite exposure, the fish were starved to increase the likelihood of copepod uptake. After another 24 h, the water in the jars was filtered to determine whether the copepod was consumed by the fish. We used family pairs in which both fish ate copepods for behavioral recordings.

### Behavior recording

To measure locomotor activity and sleep-like behavior of sticklebacks, we used fully automated cameras. The principle of this system is to place fish in experimental tanks, illuminate the tanks with infrared LEDs, and monitor fish activity throughout the day and night using infrared vision cameras (Figure 1A). We integrated four cameras into an aluminum frame to horizontally cover two 3,6 l zebrafish tanks using Techniplast (Model ZB 30). These tanks have a low depth (up to 10,5 cm), which does not allow the fish to move much along a three-dimensional axis, thereby biasing two-dimensional video tracking. Moreover, the individual tanks had a vision barrier between them, so the fish were unable to see each other during the experiment. The tanks were filled with water from the same block where the fish were kept. The oxygen supply to the experimental tanks was provided by an aquarium air pump (EHEIM). Behind each tank, we vertically placed two 850 nm LED stripes with transparent paper between the tank and LEDs for background infrared illumination.

In each behavior recording session, we transferred four exposed (infected and exposed, but not infected) and four control fish individually and randomized them into tanks. All experiments started at 11:20 AM and finished 70 h 40 min later (i.e., until 10 AM on day 3 after recording started). We tested 24 exposed and 24control fish (n=48) for activity and sleep-like behavior immediately after exposure to the parasite to detect possible effects of early infection on sleep (Figure 1B). We euthanized and dissected half of the fish (12 exposed and 12 non-exposed) with an overdose (0,5g/l) of tricaine methanesulfate (MS222) for further use immediately after the first sleep behavior recording. The other half of the fish (12 exposed and 12 non-exposed) were transferred back into the block and kept individually in net spawning boxes (JBL; 13,4 x 2,3 x 17,9 cm). We recorded this group of fish again within the same setting after 29 days of parasite exposure to test for possible long-term effects of infection on sleep (Figure 1B). Thereafter, the fish were euthanized and dissected. Temperature and light conditions (17°C, 15/9 h light/dark) did not change during parasite exposure or behavioral recording. During behavioral recordings and in the net boxes, the fish were fed once per day between 4 PM and 5 PM with frozen chironomid larvae.

### Video analysis and data processing

To track the activity and sleep-like behavior of the fish, we used the open-source Python module Phenopype (Lürig and Vidal Garcia, 2022). Within Phenopype, we drew virtual masks defining the arena for each fish, excluding the water surface, tank walls, bottom, and left or right (depending on the tank position within the system) area of the tank where bubbles emerged from the air pumps. Therefore, these masks enabled undisturbed tracking of fish. The position of the fish was tracked five times per second. All measurements were converted from from pixels to millimeters by estimating the pixel/mm ratio for each video to normalize the displacement of all fish. We then calculated the locomotor activity of each fish and estimated the displacement in mm from each frame to the next frame. After that, we calculated the sum of displacements per minute to obtain an estimate of the locomotor activity per time interval.

### Dissection and parasite screening

After recording the sleep behavior, we euthanized all fish with an overdose (0,5g/l) of tricaine methanesulfate (MS222). We measured the total and standard lengths (from the snout to the base of the caudal fin) to the nearest millimeter and weighed the fish to the nearest milligram (Table S1). We then opened the body cavity of the fish on the ventral side with sterile scissors from the urogenital pore to the gills and screened the interior for a life tapeworm. The parasite was easy to recognize at 32 dpe, but at 4 dpe, it was still very small (approximately 100 µm, Wohlleben et al., 2018). Therefore, we incubated the body and organs of the fish at room temperature in saline solution (PBS) to detach the parasite from the fish tissue. We then intensively scanned the PBS-incubated organs and bodies under a binocular with a black background and bright illumination for actively moving parasites. The brains of all 48 fish were dissected, immediately frozen in liquid nitrogen, and stored at −80°C for RNA sequencing.

### RNA extraction

Frozen brains were immediately homogenized with a cell plunger in 1 ml Ambion TRIzol reagent to avoid RNA degradation. Subsequently, the samples were sonicated in an ultrasonic bath for 10 min. After centrifugation at 4°C and 13.000 rpm for 5 min, the supernatant was transferred to a new tube, and 200 ml of chloroform was added to the brain samples and incubated for 15 min at room temperature (RT). The suspension at 10.500 rpm at 4°C for 15 min, and 400 ml of the aqueous phase was transferred to a new tube to extract RNA using the Promega SV Total RNA Isolation kit, according to the corresponding protocol. Subsequently, we eluted the RNA in 80 ml nuclease-free water and stored it at −80°C until sequencing. Four replicates were sequenced for each treatment and time point (24 samples in total). RNA quantification and qualification, library preparation, sequencing, and data analysis were performed by Biomarker Technologies (BMK) GmbH (Münster, Germany). The RNA concentration and purity were measured using a NanoDrop 2000 spectrophotometer (Thermo Fisher Scientific). RNA integrity was assessed using an RNA Nano 6000 Assay Kit on an Agilent Bioanalyzer 2100 system (Agilent Technologies, CA, USA).

### Library preparation and RNA-sequencing

A total of 1 μg of RNA per sample was used as input material for RNA sample preparation. Sequencing libraries were generated using the NEBNext UltraTM RNA Library Prep Kit for Illumina (NEB, USA), following the manufacturer’s recommendations, and index codes were added to attribute sequences to each sample. The resulting libraries were purified (AMPure XP system), and library quality was assessed using the Agilent Bioanalyzer 2100 system. Clustering of the index-coded samples was performed on a cBot Cluster Generation System using the TruSeq PE Cluster Kit v4-cBot-HS (Illumina), according to the manufacturer’s instructions. After cluster generation, the library preparations were sequenced on an Illumina Novaseq 6000 (PE150) platform and paired-end reads were generated.

#### Sequencing alignment and differential expression analysis

Adaptor sequences and low-quality reads were removed from the dataset. The raw sequences were transformed into clean reads after data processing. We obtained approximately 40–58 million clean reads per sample (Figure S3A). Between 90.64% and 91.97% of the total reads were mapped to the NCBI reference genome sequence (GAculeatus_UGA_version5). Only reads with a perfect match or one mismatch were further analyzed and annotated based on the reference genome. More than 83% of the reads were uniquely mapped to the reference genome using Hisat2 tools, resulting in a coverage of approximately 25x per sample (Sims et al., 2014). Differential expression analysis was performed using DESeq2. DESeq2 provides statistical routines for determining differential expression in digital gene expression data using a model based on negative binomial distribution (Anders and Huber, 2010). The resulting p-values were adjusted using Benjamini and Hochberg’s approach to control the false discovery rate. Genes with an adjusted P-value < 0.05 found by DESeq2 were assigned as differentially expressed.

### GO functional enrichment analysis

Gene ontology (GO) enrichment analysis of the differentially expressed genes (DEGs) was performed using the GOseq R packages based on Wallenius non-central hyper-geometric distribution, which can adjust for gene length bias in DEGs (Young et al., 2010).

### Statistical analysis of sleep-like behavior

We used the observed data on locomotor activity per minute as time series data for each fish to fit a hidden Markov model (HMM) using the hmmTMB R package (Michelot, 2022). HMMs provide a structured and probabilistic approach for modeling sequential data with hidden underlying patterns (Zucchini et al., 2017). Since sleep is strongly associated with repeated periods of behavioral inactivity (Tononi, 2000; Yokogawa et al., 2007), it can be considered a hidden pattern underlying the observed sequential data of locomotor activity (Wiggin et al., 2020). Therefore, HMMs enable the objective characterization of sleep-like behavior in organisms that are not suitable for procedures such as electroencephalographic analyses. For this purpose, we first assigned all missing observations of locomotor activity per minute to the NA to maintain the time-series structure. Overall, 4.33% of the observations were missing. Furthermore, we set all values to zero (0.0057% of the total data points) to values slightly larger than zero, to avoid introducing an additional parameter into the model. We modeled the state-dependent distributions of locomotor activity data as gamma distributions with parameterization means and standard deviations, assuming that all individuals followed the same state-dependent process.

We used an HMM with three different observation parameters (states) to model the locomotion of each fish per min. This decision was based on comparing the overlap of state-dependent distributions of two-, three-, and 4-state models and according to Pohle et al. (2017). Because of the clear differences in locomotion per minute, we interpreted state 1 as sleep-like behavior with the lowest locomotion per minute, state 2 as moderate activity with intermediate locomotion per minute, and state 3 as high activity with the highest locomotion per minute (Figure 2A, Supplementary Table 2). To investigate diel patterns in fish behavior, we modeled the state-switching probabilities as a function of the time of day by specifying trigonometric functions with wavelengths of 24 h as covariates and allowing different periodic effects in each condition (1-4 and 29-32 dpe of control, exposed, and infected fish, respectively). In addition, we included random intercepts per fish in each of the state-switching probabilities to account for the heterogeneity in behavior among individuals. However, we did not allow for any transitions between sleep-like behavior (state 1) and high activity (state 3) in our model formulation by fixing the respective parameters to zero. The diel activity/sleeping patterns in each condition were investigated at the group level by inferring the periodic stationary distribution and dwell times as a function of time of day (Koslik et al., 2023; Figures 2E and 2F).

Decoding the most probable underlying state sequence for each individual using the Viterbi algorithm revealed that overall, the fish spent 18.7% in state 1, 59.1% in state 2, and 22.2% in state 3. The marginal distribution of the fitted HMM accurately captures the underlying empirical distribution (Figure 2A). To further assess the goodness-of-fit, we simulated the data from the fitted HMM and calculated pseudo-residuals (Figure S1). Although the model checks revealed a slight lack of fit regarding the tails of the distribution and the observed autocorrelation, the overall model fit was satisfactory.

## Notes

### Competing Interest Statement

The authors have declared no competing interest.

## References

Anders S, Huber W. 2010. Differential expression analysis for sequence count data. Nature Precedings 2010 1–1. doi:10.1038/npre.2010.4282.1

Baer J, Gugele SM, Roch S, Brinker A. 2022. Stickleback mass occurrence driven by spatially uneven parasite pressure? Insights into infection dynamics, host mortality, and epizootic variability. Parasitol Res 121:1607–1619. doi:10.1007/S00436-022-07517-4

Barber I, Scharsack JP. 2010. The three-spined stickleback-Schistocephalus solidus system: An experimental model for investigating host-parasite interactions in fish. Parasitology. doi:10.1017/S0031182009991466

Barber I, Svensson PA. 2003. Effects of experimental Schistocephalus solidus infections on growth, morphology and sexual development of female three-spined sticklebacks, Gasterosteus aculeatus. Parasitology 126:359–367. doi:10.1017/S0031182002002925

Barber I, Walker P, Svensson PA. 2004. Behavioural Responses to Simulated Avian Predation in Female Three Spined Sticklebacks: The Effect of Experimental Schistocephalus Solidus Infections. undefined 141:1425–1440. doi:10.1163/1568539042948231

Bédard T, Mountney C, Kent P, Anisman H, Merali Z. 2007. Role of gastrin-releasing peptide and neuromedin B in anxiety and fear-related behavior. Behavioural Brain Research 179:133–140. doi:10.1016/j.bbr.2007.01.021

Benesh DP, Hafer N. 2012. Growth and ontogeny of the tapeworm Schistocephalus solidus in its copepod first host affects performance in its stickleback second intermediate host. Parasit Vectors 5:1–10. doi:10.1186/1756-3305-5-90

Besedovsky L, Lange T, Haack M. 2019. The sleep-immune crosstalk in health and disease. Physiol Rev. doi:10.1152/physrev.00010.2018

Borbély AA, Daan S, Wirz-Justice A, Deboer T. 2016. The two-process model of sleep regulation: A reappraisal. J Sleep Res 25:131–143. doi:10.1111/JSR.12371

Brochu M-P, Aubin-Horth N. 2021. Shedding light on the circadian clock of the threespine stickleback. Journal of Experimental Biology 224. doi:10.1242/JEB.242970

Brown RE, Basheer R, McKenna JT, Strecker RE, McCarley RW. 2012. Control of sleep and wakefulness. Physiol Rev 92:1087–1187. doi:10.1152/physrev.00032.2011

Brown TM, Hughes AT, Piggins HD. 2005. Gastrin-Releasing Peptide Promotes Suprachiasmatic Nuclei Cellular Rhythmicity in the Absence of Vasoactive Intestinal Polypeptide-VPAC2 Receptor Signaling. Journal of Neuroscience 25:11155–11164. doi:10.1523/JNEUROSCI.3821-05.2005

Campbell SS, Tobler I. 1984. Animal sleep: A review of sleep duration across phylogeny. Neurosci Biobehav Rev. doi:10.1016/0149-7634(84)90054-X

Chacón PJ, del Marco Á, Arévalo Á, Domínguez-Giménez P, García-Segura LM, Rodríguez-Tébar A. 2015. Cerebellin 4, a synaptic protein, enhances inhibitory activity and resistance of neurons to amyloid-β toxicity. Neurobiol Aging 36:1057–1071. doi:10.1016/J.NEUROBIOLAGING.2014.11.006

Chang MX, Xiong F, Wu XM, Hu YW. 2021. The expanding and function of NLRC3 or NLRC3-like in teleost fish: Recent advances and novel insights. Dev Comp Immunol 114. doi:10.1016/J.DCI.2020.103859

Chen F, Lu XJ, Nie L, Ning YJ, Chen J. 2018. Molecular characterization of a CC motif chemokine 19-like gene in ayu (Plecoglossus altivelis) and its role in leukocyte trafficking. Fish Shellfish Immunol 72:301–308. doi:10.1016/J.FSI.2017.11.012

Chitu V, ölen Gokhan S, Nandi S, Mehler MF, Stanley ER. 2016. Emerging Roles for CSF-1 Receptor and its Ligands in the Nervous System. doi:10.1016/j.tins.2016.03.005

Demandt N, Praetz M, Kurvers RHJM, Krause J, Kurtz J, Scharsack JP. 2020. Parasite infection disrupts escape behaviours in fish shoals. Proceedings of the Royal Society B 287. doi:10.1098/RSPB.2020.1158

Drake CL, Roehrs TA, Royer H, Koshorek G, Turner RB, Roth T. 2000. Effects of an experimentally induced rhinovirus cold on sleep, performance, and daytime alertness. Physiol Behav 71:75–81. doi:10.1016/S0031-9384(00)00322-X

Fang J, Sanborn CK, Renegar KB, Krueger JM, Majde JA. 2016. Influenza Viral Infections Enhance Sleep in Mice: 103181/00379727-210-43945 210:242–252. doi:10.3181/00379727-210-43945

Goldstein N, Levine BJ, Loy KA, Duke WL, Meyerson OS, Jamnik AA, Carter ME. 2018. Hypothalamic Neurons that Regulate Feeding Can Influence Sleep/Wake States Based on Homeostatic Need. Curr Biol 28:3736–3747.e3. doi:10.1016/J.CUB.2018.09.055

Grecias L, Hebert FO, Alves VA, Barber I, Aubin-Horth N. 2020. Host behaviour alteration by its parasite: from brain gene expression to functional test. Proceedings of the Royal Society B 287. doi:10.1098/RSPB.2020.2252

Guo F, He H, Fu ZC, Huang S, Chen T, Papasian CJ, Morse LR, Xu Y, Battaglino RA, Yang XF, Jiang Z, Xin HB, Fu M. 2015. Adipocyte-derived PAMM suppresses macrophage inflammation by inhibiting MAPK signalling. Biochem J 472:309–318. doi:10.1042/BJ20150019

Hahn MA, Piecyk A, Jorge F, Cerrato R, Kalbe M, Dheilly NM. 2022. Host phenotype and microbiome vary with infection status, parasite genotype, and parasite microbiome composition. Mol Ecol 31:1577–1594. doi:10.1111/MEC.16344

Hammerschmidt K, Kurtz J. 2005. Surface carbohydrate composition of a tapeworm in its consecutive intermediate hosts: Individual variation and fitness consequences. Int J Parasitol 35:1499–1507. doi:10.1016/J.IJPARA.2005.08.011

Hammerschmidt K, Kurtz J. 2009. Chapter 5 Ecological Immunology of a Tapeworms’ Interaction with its Two Consecutive Hosts. Adv Parasitol 68:111–137. doi:10.1016/S0065-308X(08)00605-2

Hanington PC, Belosevic M. 2007. Interleukin-6 family cytokine M17 induces differentiation and nitric oxide response of goldfish (Carassius auratus L.) macrophages. Dev Comp Immunol 31:817–829. doi:10.1016/J.DCI.2006.12.001

Hayashi F, Smith KD, Ozinsky A, Hawn TR, Yi EC, Goodlett DR, Eng JK, Akira S, Underhill DM, Aderem A. 2001. The innate immune response to bacterial flagellin is mediated by Toll-like receptor 5. Nature 2001 410:6832 410:1099–1103. doi:10.1038/35074106

He Y, Jones CR, Fujiki N, Xu Y, Guo B, Holder JL, Rossner MJ, Nishino S, Fu YH. 2009. The Transcriptional Repressor DEC2 Regulates Sleep Length in Mammals. Science 325:866. doi:10.1126/SCIENCE.1174443

Hu J, Yang Z, Li X, Lu H. 2016. C-C motif chemokine ligand 20 regulates neuroinflammation following spinal cord injury via Th17 cell recruitment. J Neuroinflammation 13:1–14. doi:10.1186/S12974-016-0630-7/FIGURES/9

Im E, Jung J, Rhee SH. 2012. Toll-Like Receptor 5 Engagement Induces Interleukin-17C Expression in Intestinal Epithelial Cells. Journal of Interferon & Cytokine Research 32:583. doi:10.1089/JIR.2012.0053

Imeri L, Opp MR. 2009. How (and why) the immune system makes us sleep. Nature Reviews Neuroscience 2009 10:3 10:199–210. doi:10.1038/nrn2576

Ito T, Carson WF, Cavassani KA, Connett JM, Kunkel SL. 2011. CCR6 as a mediator of immunity in the lung and gut. Exp Cell Res 317:613–619. doi:10.1016/J.YEXCR.2010.12.018

Jiang H, Tsang L, Wang H, Liu C. 2021. IFI44L as a Forward Regulator Enhancing Host Antituberculosis Responses. J Immunol Res 2021. doi:10.1155/2021/5599408

Kagawa Y, Low YL, Pyun J, Doglione U, Short JL, Pan Y, Nicolazzo JA. 2023. Fatty Acid-Binding Protein 4 is Essential for the Inflammatory and Metabolic Response of Microglia to Lipopolysaccharide. J Neuroimmune Pharmacol 18:448–461. doi:10.1007/S11481-023-10079-6

Kimura-Takeuchi M, Majde JA, Toth LA, Krueger JM. 1992. Influenza virus-induced changes in rabbit sleep and acute phase responses. 101152/ajpregu19922635R1115 **263**. doi:10.1152/AJPREGU.1992.263.5.R1115

Koslik J-O, Feldmann CC, Mews S, Michels R, Langrock R. 2023. Inference on the state process of periodically inhomogeneous hidden Markov models for animal behavior. doi: 10.48550/arXiv.2312.14583

Krueger JM, Frank MG, Wisor JP, Roy S. 2016. Sleep function: Toward elucidating an enigma. Sleep Med Rev 28:46–54. doi:10.1016/J.SMRV.2015.08.005

Kubota T, Brown RA, Fang J, Krueger JM. 2001a. Interleukin-15 and interleukin-2 enhance non-REM sleep in rabbits. Am J Physiol Regul Integr Comp Physiol 281:1004–1012. doi:10.1152/AJPREGU.2001.281.3.R1004

Kubota T, Fang J, Brown RA, Krueger JM. 2001b. Interleukin-18 promotes sleep in rabbits and rats. Am J Physiol Regul Integr Comp Physiol 281:828–838. doi:10.1152/AJPREGU.2001.281.3.R828

Lazarus M, Oishi Y, Bjorness TE, Greene RW. 2019. Gating and the need for sleep: Dissociable effects of adenosine a1and a2areceptors. Front Neurosci 13:463244. doi:10.3389/FNINS.2019.00740/BIBTEX

Lee JM, Kim H, Baek SH. 2021. Unraveling the physiological roles of retinoic acid receptor-related orphan receptor α. Experimental & Molecular Medicine 2021 53:9 53:1278–1286. doi:10.1038/s12276-021-00679-8

Lesku JA, Schmidt MH. 2022. Energetic costs and benefits of sleep. Current Biology 32:R656–R661. doi:10.1016/j.cub.2022.04.004

Liu X, Cao X, Wang S, Ji G, Zhang S, Li H. 2017. Identification of Ly2 members as antimicrobial peptides from zebrafish Danio rerio. Biosci Rep 37. doi:10.1042/BSR20160265

Lürig MD, Vidal Garcia M. 2022. phenopype: A phenotyping pipeline for Python. Methods Ecol Evol 13:569–576. doi:10.1111/2041-210X.13771

Lye E, Mirtsos C, Suzuki N, Suzuki S, Yeh WC. 2004. The role of interleukin 1 receptor-associated kinase-4 (IRAK-4) kinase activity in IRAK-4-mediated signaling. J Biol Chem 279:40653–40658. doi:10.1074/JBC.M402666200

McAlpine CS, Kiss MG, Rattik S, He S, Vassalli A, Valet C, Anzai A, Chan CT, Mindur JE, Kahles F, Poller WC, Frodermann V, Fenn AM, Gregory AF, Halle L, Iwamoto Y, Hoyer FF, Binder CJ, Libby P, Tafti M, Scammell TE, Nahrendorf M, Swirski FK. 2019. Sleep modulates haematopoiesis and protects against atherosclerosis. Nature. doi:10.1038/s41586-019-0948-2

Michelot T. 2022. hmmTMB: Hidden Markov models with flexible covariate effects in R. arXiv:2211.14139. doi: 10.48550/arXiv.2211.14139

Molbert N, Goutte A. 2022. Narrower isotopic niche size in fish infected by the intestinal parasite Pomphorhynchus sp. compared to uninfected ones. J Fish Biol 101:1466–1473. doi:10.1111/JFB.15217

Monti JM, Jantos H. 2008. Activation of the serotonin 5-HT3 receptor in the dorsal raphe nucleus suppresses REM sleep in the rat. Prog Neuropsychopharmacol Biol Psychiatry 32:940–947. doi:10.1016/J.PNPBP.2007.12.024

Mullington J, Korth C, Hermann DM, Orth A, Galanos C, Holsboer F, Pollmächer T. 2000. Dose-dependent effects of endotoxin on human sleep. Am J Physiol Regul Integr Comp Physiol 278. doi:10.1152/AJPREGU.2000.278.4.R947

Nebert DW, Vasiliou V. 2004. Analysis of the glutathione S-transferase (GST) gene family. Hum Genomics 1:460–464. doi:10.1186/1479-7364-1-6-460

Nordeide JT, Matos F. 2016. Solo Schistocephalus solidus tapeworms are nasty. Parasitology 143:1301–1309. doi:10.1017/S0031182016000676

Opp MR. 2005. Cytokines and sleep. Sleep Med Rev 9:355–364. doi:10.1016/J.SMRV.2005.01.002

Panula P, Aarnisalo AA, Wasowicz K. 1996. Neuropeptide FF, a mammalian neuropeptide with multiple functions. Prog Neurobiol 48:461–479. doi:10.1016/0301-0082(96)00001-9

Patel AK, Reddy V, Araujo JF. 2021. Physiology, Sleep Stages. StatPearls.

Pellegrino R, Kavakli IH, Goel N, Cardinale CJ, Dinges DF, Kuna ST, Maislin G, Van Dongen HPA, Tufik S, Hogenesch JB, Hakonarson H, Pack AI. 2014. A novel BHLHE41 variant is associated with short sleep and resistance to sleep deprivation in humans. Sleep 37:1327–1336. doi:10.5665/SLEEP.3924

Piecyk A, Ritter M, Kalbe M. 2019. The right response at the right time: Exploring helminth immune modulation in sticklebacks by experimental coinfection. Mol Ecol 28:2668–2680. doi:10.1111/MEC.15106

Pilla-Moffett D, Barber MF, Taylor GA, Coers J. 2016. Interferon-inducible GTPases in host resistance, inflammation and disease. J Mol Biol 428:3495. doi:10.1016/J.JMB.2016.04.032

Pohle J, Langrock R, van Beest FM, Schmidt NM. 2017. Selecting the Number of States in Hidden Markov Models: Pragmatic Solutions Illustrated Using Animal Movement. J Agric Biol Environ Stat 22:270–293. doi:10.1007/S13253-017-0283-8/TABLES/2

Preston BT, Capellini I, Mcnamara P, Barton RA, Nunn CL. 2009. Parasite resistance and the adaptive significance of sleep. doi:10.1186/1471-2148-9-7

Quinn TP, Kendall NW, Rich HB, Chasco BE. 2012. Diel vertical movements, and effects of infection by the cestode Schistocephalus solidus on daytime proximity of three-spined sticklebacks Gasterosteus aculeatus to the surface of a large Alaskan lake. Oecologia 168:43–51. doi:10.1007/S00442-011-2071-4

Rattenborg NC, De La Iglesia HO, Kempenaers B, Lesku JA, Meerlo P, Scriba MF. 2017. Sleep research goes wild: new methods and approaches to investigate the ecology, evolution and functions of sleep. Philos Trans R Soc Lond B Biol Sci 372. doi:10.1098/RSTB.2016.0251

Rebeaud F, Hailfinger S, Posevitz-Fejfar A, Tapernoux M, Moser R, Rueda D, Gaide O, Guzzardi M, Iancu EM, Rufer N, Fasel N, Thome M. 2008. The proteolytic activity of the paracaspase MALT1 is key in T cell activation. Nature Immunology 2008 9:3 9:272–281. doi:10.1038/ni1568

Schärer L, Wedekind C. 1999. Lifetime reproductive output in a hermaphrodite cestode when reproducing alone or pairs: A time cost of pairing. Evol Ecol 13:381–394. doi:10.1023/A:1006789110502

Scharsack JP, Koch K, Hammerschmidt K. 2007. Who is in control of the stickleback immune system: interactions between Schistocephalus solidus and its specific vertebrate host. Proceedings of the Royal Society B: Biological Sciences 274:3151–3158. doi:10.1098/RSPB.2007.1148

Scharsack JP, Wieczorek B, Schmidt-Drewello A, Büscher J, Franke F, Moore A, Branca A, Witten A, Stoll M, Bornberg-Bauer E, Wicke S, Kurtz J. 2021. Climate change facilitates a parasite’s host exploitation via temperature-mediated immunometabolic processes. Glob Chang Biol 27:94–107. doi:10.1111/GCB.15402

Schmid-Hempel P. 2008. Immune defence, parasite evasion strategies and their relevance for macroscopic phenomena such as virulence. Philosophical Transactions of the Royal Society B: Biological Sciences 364:85–98. doi:10.1098/RSTB.2008.0157

Schmidt MH. 2014. The energy allocation function of sleep: A unifying theory of sleep, torpor, and continuous wakefulness. Neurosci Biobehav Rev 47:122–153. doi:10.1016/J.NEUBIOREV.2014.08.001

Sevenster P, Feuth-De Bruijn E, Huisman JJ. 1995. Temporal Structure in Stickleback Behaviour. Behaviour 132:1267–1284. doi:10.1163/156853995X00577

Sher S, Green A, Khatib S, Dagan Y. 2021. The Possible Role of Endozepines in Sleep Regulation and Biomarker of Process S of the Borbély Sleep Model. Chronobiol Int 38:122–128. doi:10.1080/07420528.2020.1849252

Sims D, Sudbery I, Ilott NE, Heger A, Ponting CP. 2014. Sequencing depth and coverage: key considerations in genomic analyses. Nature Reviews Genetics 2014 15:2 15:121–132. doi:10.1038/nrg3642

Staner L, Linker T, Toussaint M, Danjou P, Roegel JC, Luthringer R, Le Fur G, Macher JP. 2001. Effects of the selective activation of 5-HT3 receptors on sleep: a polysomnographic study in healthy volunteers. European Neuropsychopharmacology 11:301–305. doi:10.1016/S0924-977X(01)00099-2

Stumbo AD, Poulin R. 2016. Possible mechanism of host manipulation resulting from a diel behaviour pattern of eye-dwelling parasites? Parasitology 143:1261–1267. doi:10.1017/S0031182016000810

Sun YL, Zhang XY, He N, Sun T, Zhuang Y, Fang Q, Wang KR, Wang R. 2012. Neuropeptide FF activates ERK and NF kappa B signal pathways in differentiated SH-SY5Y cells. Peptides (NY) 38:110–117. doi:10.1016/J.PEPTIDES.2012.08.019

Tai H-H, Cho H, Tong M, Ding Y. 2006. NAD+-Linked 15-Hydroxyprostaglandin Dehydrogenase: Structure and Biological Functions. Curr Pharm Des 12:955–962. doi:10.2174/138161206776055958

Talarico M, Seifert F, Lange J, Sachser N, Kurtz J, Scharsack JP. 2017. Specific manipulation or systemic impairment? Behavioural changes of three-spined sticklebacks (Gasterosteus aculeatus) infected with the tapeworm Schistocephalus solidus. Behav Ecol Sociobiol 71:1–10. doi:10.1007/S00265-017-2265-9

Tononi G. 2000. Correlates of sleep and waking in Drosophila melanogaster. Science (1979). doi:10.1126/science.287.5459.1834

Toth LA. 2019. Microbial Modulation of Arousal. Handbook of Behavioral State Control 641–657. doi:10.1201/9780429114373-40

Toth LA, Kreuger JM. 1988. Alteration of sleep in rabbits by Staphylococcus aureus infection. Infect Immun. doi:10.1128/iai.56.7.1785-1791.1988

Trappes R, Nematipour B, Kaiser MI, Krohs U, Van Benthem KJ, Ernst UR, Gadau J, Korsten P, Kurtz J, Schielzeth H, Schmoll T, Takola E. 2022. How Individualized Niches Arise: Defining Mechanisms of Niche Construction, Niche Choice, and Niche Conformance. Bioscience 72:538–548. doi:10.1093/BIOSCI/BIAC023

Van Den Pol AN, Yao Y, Fu LY, Foo K, Huang H, Coppari R, Lowell BB, Broberger C. 2009. Neuromedin B and Gastrin-Releasing Peptide Excite Arcuate Nucleus Neuropeptide Y Neurons in a Novel Transgenic Mouse Expressing Strong Renilla Green Fluorescent Protein in NPY Neurons. The Journal of Neuroscience 29:4622. doi:10.1523/JNEUROSCI.3249-08.2009

Wang X, Yin G, Zhang W, Song K, Zhang L, Guo Z. 2021. Prostaglandin Reductase 1 as a Potential Therapeutic Target for Cancer Therapy. Front Pharmacol 12. doi:10.3389/FPHAR.2021.717730

Weber JN, Steinel NC, Peng F, Shim KC, Lohman BK, Fuess LE, Subramanian S, De Lisle SP, Bolnick DI. 2022. Evolutionary gain and loss of a pathological immune response to parasitism. Science (1979) 377:1206–1211. doi:10.1126/science.abo3411

Wiggin TD, Goodwin PR, Donelson NC, Liu C, Trinh K, Sanyal S, Griffith LC. 2020. Covert sleep-related biological processes are revealed by probabilistic analysis in Drosophila. Proc Natl Acad Sci U S A 117:10024–10034. doi:10.1073/PNAS.1917573117

Wilcox AG, Vizor L, Parsons MJ, Banks G, Nolan PM. 2017. Inducible Knockout of Mouse Zfhx3 Emphasizes Its Key Role in Setting the Pace and Amplitude of the Adult Circadian Clock. J Biol Rhythms 32:433–443. doi:10.1177/0748730417722631

Wohlleben AM, Franke F, Hamley M, Kurtz J, Scharsack JP. 2018. Early stages of infection of three-spined stickleback (Gasterosteus aculeatus) with the cestode Schistocephalus solidus. J Fish Dis 41:1701–1708. doi:10.1111/JFD.12876

Yan A, Zhang T, Yang X, Shao J, Fu N, Shen F, Fu Y, Xia W. 2016. Thromboxane A2 receptor antagonist SQ29548 reduces ischemic stroke-induced microglia/macrophages activation and enrichment, and ameliorates brain injury. Scientific Reports 2016 6:1 6:1–13. doi:10.1038/srep35885

Yan Y, Su J, Zhang Z. 2022. The CXCL12/CXCR4/ACKR3 Response Axis in Chronic Neurodegenerative Disorders of the Central Nervous System: Therapeutic Target and Biomarker. Cell Mol Neurobiol 42:2147–2156. doi:10.1007/S10571-021-01115-1

Yo B, Haubrock PJ, Aguirre W, Hudson CM, Lucek K, Marques DA, Alexander TJ, Moosmann M, Spaak P, Seehausen O, Matthews B. 2021. Threespine Stickleback in Lake Constance: The Ecology and Genomic Substrate of a Recent Invasion. Frontiers in Ecology and Evolution | www.frontiersin.org 1:611672. doi:10.3389/fevo.2020.611672

Yokogawa T, Marin W, Faraco J, Pézeron G, Appelbaum L, Zhang J, Rosa F, Mourrain P, Mignot E. 2007. Characterization of sleep in zebrafish and insomnia in hypocretin receptor mutants. PLoS Biol. doi:10.1371/journal.pbio.0050277

Yoshizawa M, Robinson BG, Duboué ER, Masek P, Jaggard JB, O’Quin KE, Borowsky RL, Jeffery WR, Keene AC. 2015. Distinct genetic architecture underlies the emergence of sleep loss and prey-seeking behavior in the Mexican cavefish. BMC Biol 13:1–12. doi:10.1186/S12915-015-0119-3

Young MD, Wakefield MJ, Smyth GK, Oshlack A. 2010. Gene ontology analysis for RNA-seq: accounting for selection bias. Genome Biol 11:1–12. doi:10.1186/GB-2010-11-2-R14

Zhong Y, Ye Q, Chen C, Wang M, Wang H. 2018. Ezh2 promotes clock function and hematopoiesis independent of histone methyltransferase activity in zebrafish. Nucleic Acids Res 46:3382. doi:10.1093/NAR/GKY101

Zucchini W, Macdonald IL, Langrock R. 2017. Hidden Markov models for time series: An introduction using R, second edition. Hidden Markov Models for Time Series: An Introduction Using R, Second Edition 1–370. doi:10.1007/s13252-016-0273-2

