## Supplementary Tables & Figures for "Tapeworm infection affects sleep behavior in three-spined stickleback"

### Supplemental information

#### Supplementary tables:

**Supplementary Table 1:** Fish data with information about parasite exposure, sex, total and standard length, weight, infection status, and parasite mass. Total length was measured from the snout to the end of the caudal fin and standard length (in brackets) was measured from the snout to the caudal peduncle. Missing values are indicated with NA.

| Fish ID | Parasite exposed? | Sex | Total (standard) length [cm] | Weight [g] | Infected? | Parasite mass [mg] |
|---|---|---|---|---|---|---|
| 1 | yes | f | 5.2 (4.1) | 1.008 | no | - |
| 2 | no | m | 5.5 (4.9) | 1.205 | - | - |
| 3 | no | f | 4.9 (4.4) | 0.894 | no | - |
| 4 | yes | m | 4.7 (4.2) | 0.753 | yes | NA |
| 5 | yes | m | 5.2 (4.2) | 0.932 | no | - |
| 6 | no | m | 4.7 (4.2) | 0.705 | - | - |
| 7 | yes | f | 5.4 (4.7) | 1.188 | no | - |
| 8 | no | f | 5.5 (4.9) | 1.179 |  | - |
| 9 | yes | f | 5.4 (4.8) | 1.12 | yes | 18 |
| 10 | no | f | 5.3 (4.7) | 1.041 | - | - |
| 11 | yes | m | 5.6 (5.0) | 1.137 | no | - |
| 12 | no | m | 5.4 (4.8) | 1.099 | - | - |
| 13 | yes | f | 5.6 (5.0) | 1.252 | no | - |
| 14 | no | m | 5.6 (5.0) | 1.285 | - | - |
| 15 | yes | m | 4.9 (4.3) | 0.802 | no | - |
| 16 | no | m | 5.5 (4.8) | 1.066 | - | - |
| 17 | yes | f | 5.4 (4.8) | 1.109 | no | - |
| 18 | no | f | 4.8 (4.4) | 0.82 | - | - |
| 19 | yes | f | 5.3 (4.7) | 1.145 | yes | 10 |
| 20 | no | m | 5.0 (4.4) | 0.875 | - | - |
| 21 | yes | m | 5.3 (4.7) | 1.042 | yes | 19 |
| 22 | no | m | 4.9 (4.3) | 0.836 | - | - |
| 23 | yes | f | 5.4 (4.8) | 1.16 | yes | 8 |
| 24 | no | f | 5.6 (4.9) | 1.202 | - | - |
| 25 | yes | m | 5.2 (NA) | 1.002 | yes | NA |
| 26 | no | f | 5.4 (NA) | 1.166 | - | - |
| 27 | yes | m | 5.3 (NA) | 1.029 | yes | NA |
| 28 | no | m | 5 (NA) | 0.934 | - | - |
| 29 | yes | f | 5.6 (NA) | 1.224 | no | - |
| 30 | no | f | 5.4 (NA) | 1.109 | - | - |
| 31 | yes | f | 5.7 (NA) | 1.211 | no | - |
| 32 | no | f | 5.1 (NA | 0.936 | - | - |
| 33 | yes | f | 5.7 (5.2) | 1.426 | no | - |
| 34 | no | f | 5.9 (5.3) | 1.373 | - | - |
| 35 | yes | m | 6.4 (5.4) | 1.832 | no | - |
| 36 | no | f | 5.5 (4.9) | 1.019 | - | - |
| 37 | yes | f | 5.5 (4.8) | 1.086 | no | - |
| 38 | no | f | 5.8 (5.2) | 1.424 | - | - |
| 39 | yes | f | 5.8 (5.1) | 1.399 | no | - |
| 40 | no | m | 5.2 (4.6) | 0.999 | - | - |
| 41 | yes | m | 6.1 (5.4) | 1.677 | yes | NA |
| 42 | no | f | 5.9 (5.2) | 1.371 | - | - |
| 43 | yes | f | 6.4 (5.7) | 1.767 | no | - |
| 44 | no | m | 5.8 (5.2) | 1.361 | - | - |
| 45 | yes | f | 5.7 (5.1) | 1.358 | no | - |
| 46 | no | f | 5.7 (5.1) | 1.274 | - | - |
| 47 | yes | f | 5.5 (4.9) | 1.196 | yes | NA |
| 48 | no | m | 5.4 (4.8) | 1.103 | - | - |

**Supplementary table 2:** Mean and standard deviation (SD) of the observation parameters (states) estimated by the HMM.

| | State 1 | State 2 | State 3 |
|---|---|---|---|
| Mean locomotor activity (m/min) | 0.304 | 0.987 | 2.703 |
| SD locomotor activity (m/min) | 0.297 | 0.773 | 1.954 |

#### Supplementary Figures

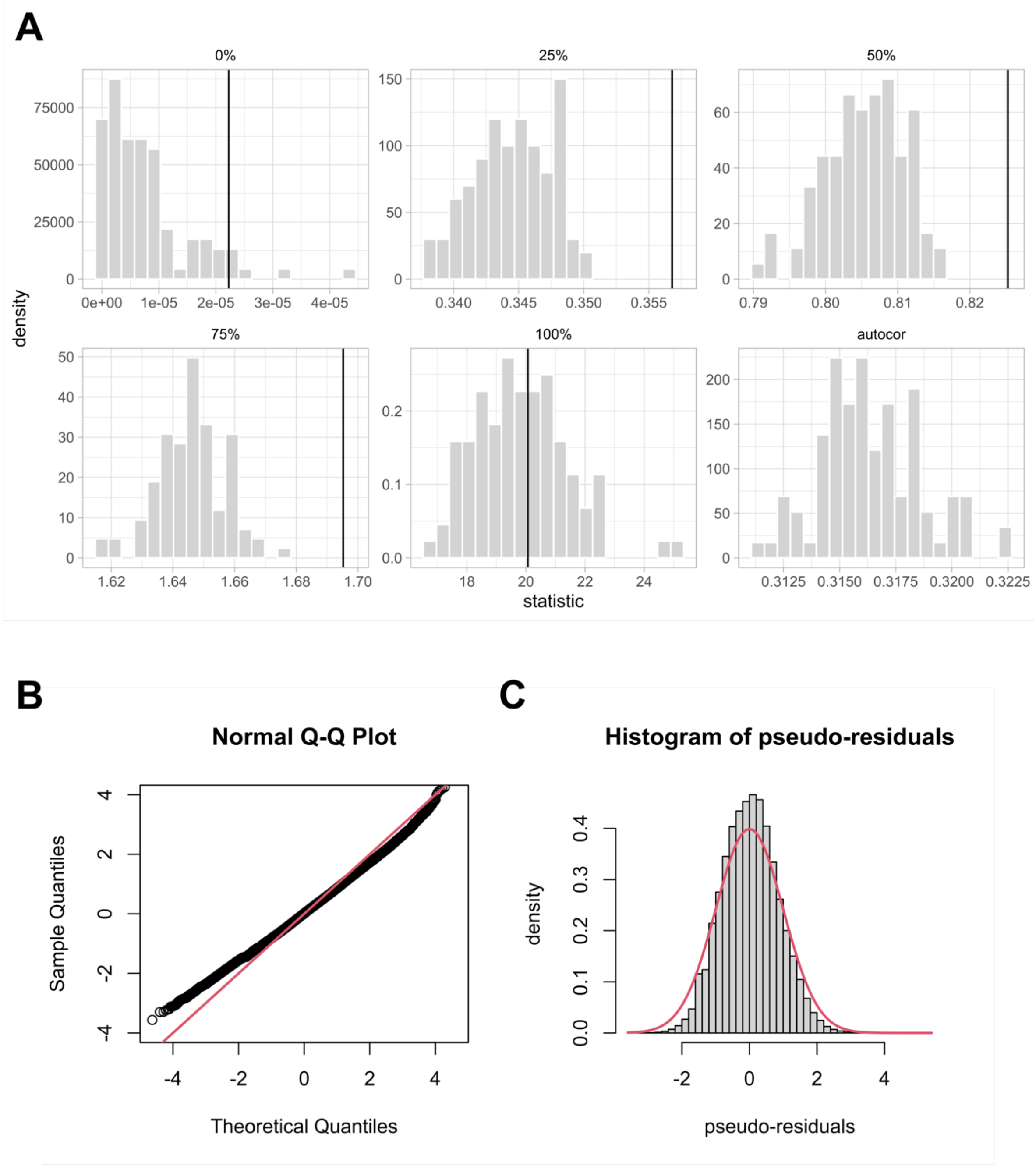


**Supplementary Figure 1:** Assessment of goodness-of-model-fit. **(A)** histograms of minimum, maximum, median, and quartiles of simulated observations under the fitted HMM and autocorrelation between consecutive observations based on 100 simulated data sets. The vertical line represents the value observed in the real data. The observed value for the autocorrelation is 0.395 and therefore exceeds the scale of the simulated autocorrelation histogram. **(B)** Normality of pseudo-residuals under the fitted HMM. Sample quantiles (y-axis)

are plotted against theoretical quantiles (x-axis) and the red line corresponds to the standard normal distribution. **(C)** Histogram of pseudo-residuals with the red line corresponding to the standard normal distribution.

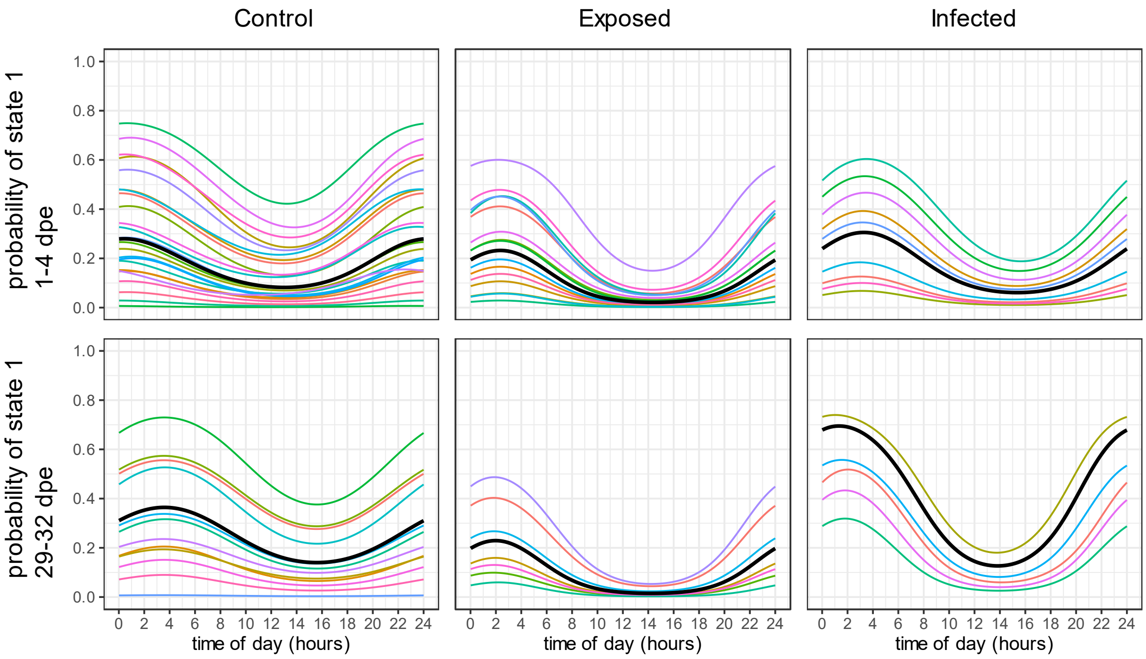


**Supplementary Figure 2:** Heterogeneity in sleep behavior among individuals. Each colored line represents the probability of an individual occupying the sleep state 1, corresponding to the periodic stationary distribution of the HMM, per recording time. The black line indicates the mean for each treatment and recording time.

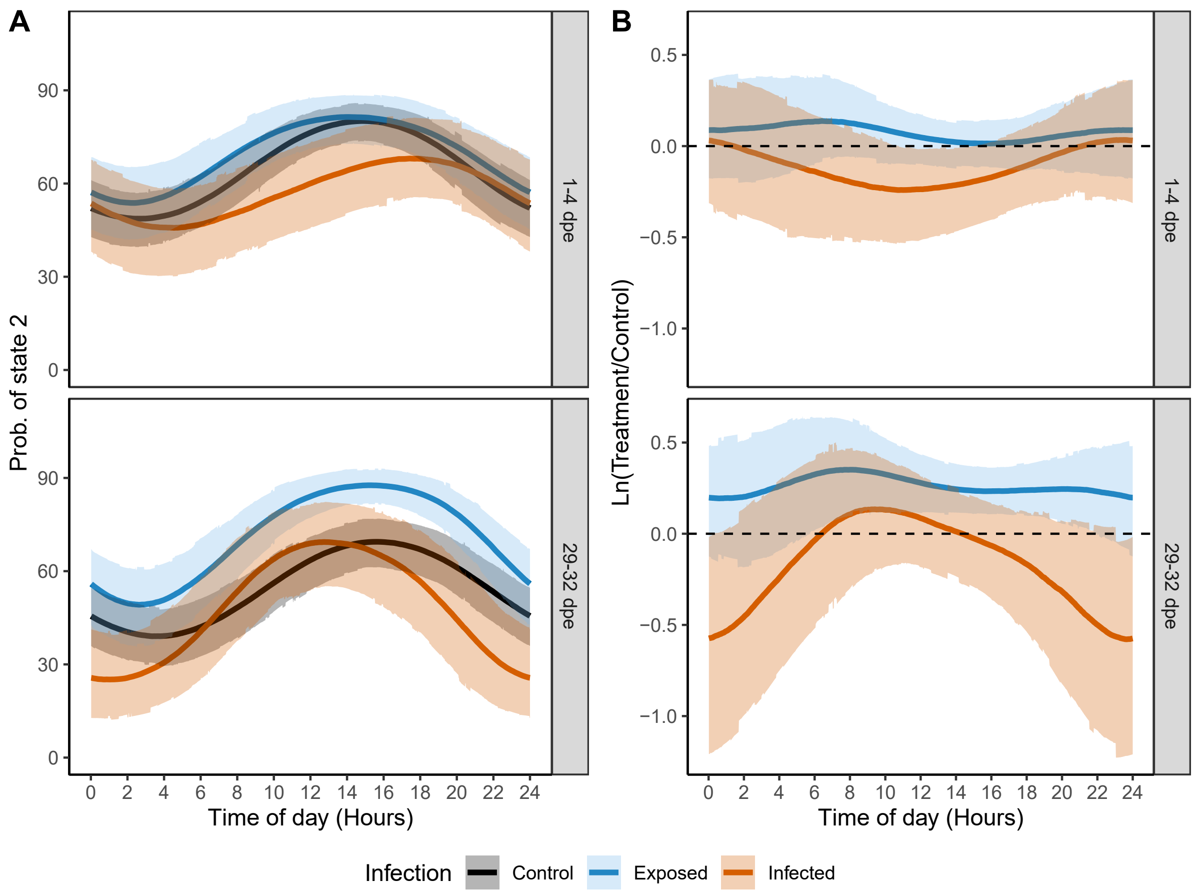


**Supplementary Figure 3. (A)** Probabilities (i.e., expected percentages) for *Control*, *Exposed*, and *Infected* fish of occupying state 2 (moderate activity), corresponding to the periodic stationary distribution of the HMM, per recording time (1-4- and 29-32 dpe). Middle lines display the mean probabilities and upper and lower areas the respective 95% confidence intervals. **(B)** Logarithmic ratio of the deviation in state 2 of exposed and infected fish from the respective control (dashed line) derived from simulations based on the HMM. Middle lines display the means and upper and lower areas the 95% confidence intervals of the distribution.

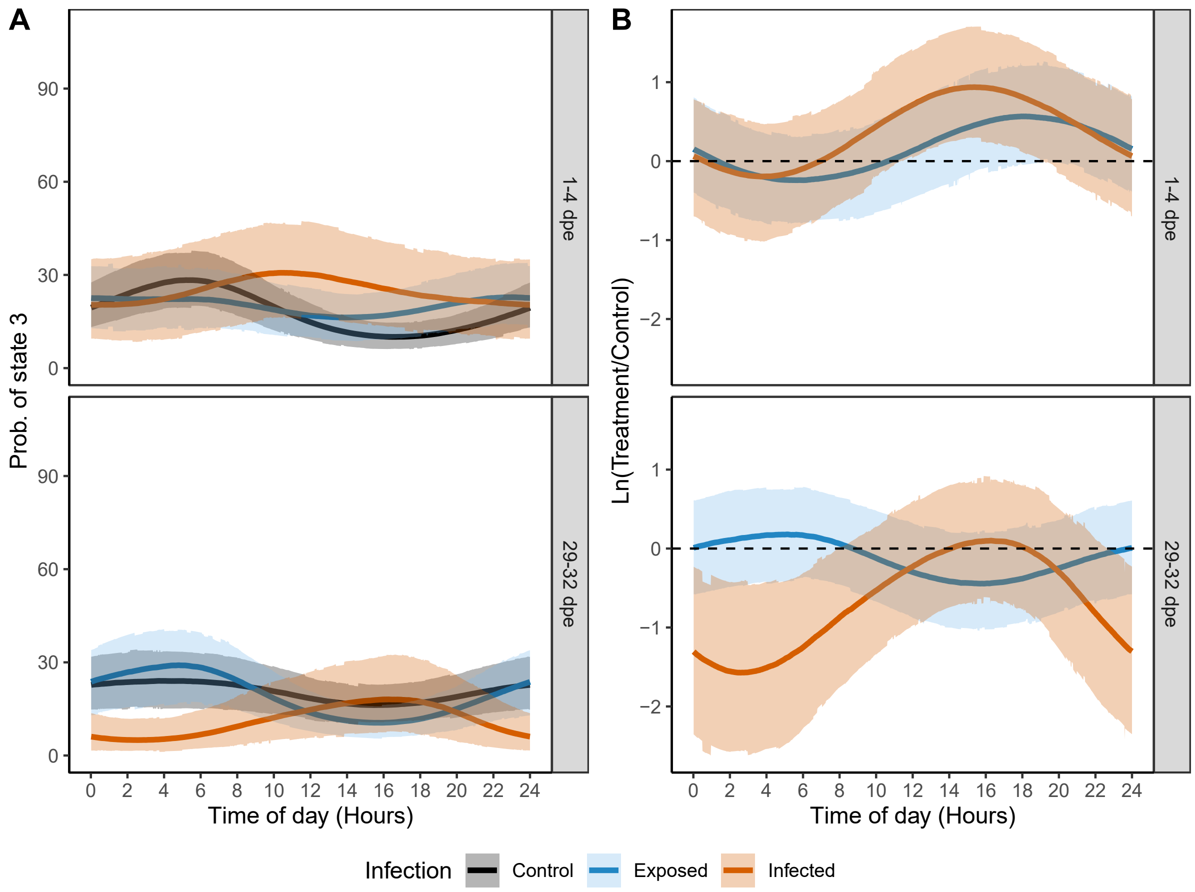


**Supplementary Figure 4. (A)** Probabilities (i.e., expected percentages) for *Control*, *Exposed*, and *Infected* fish of occupying state 3 (high activity), corresponding to the periodic stationary distribution of the HMM, per recording time (1-4- and 29-32 dpe). Middle lines display the mean probabilities and upper and lower areas the respective 95% confidence intervals. **(B)** Logarithmic ratio of the deviation in state 3 of exposed and infected fish from the respective control (dashed line) derived from simulations based on the HMM. Middle lines display the means and upper and lower areas the 95% confidence intervals of the distribution.

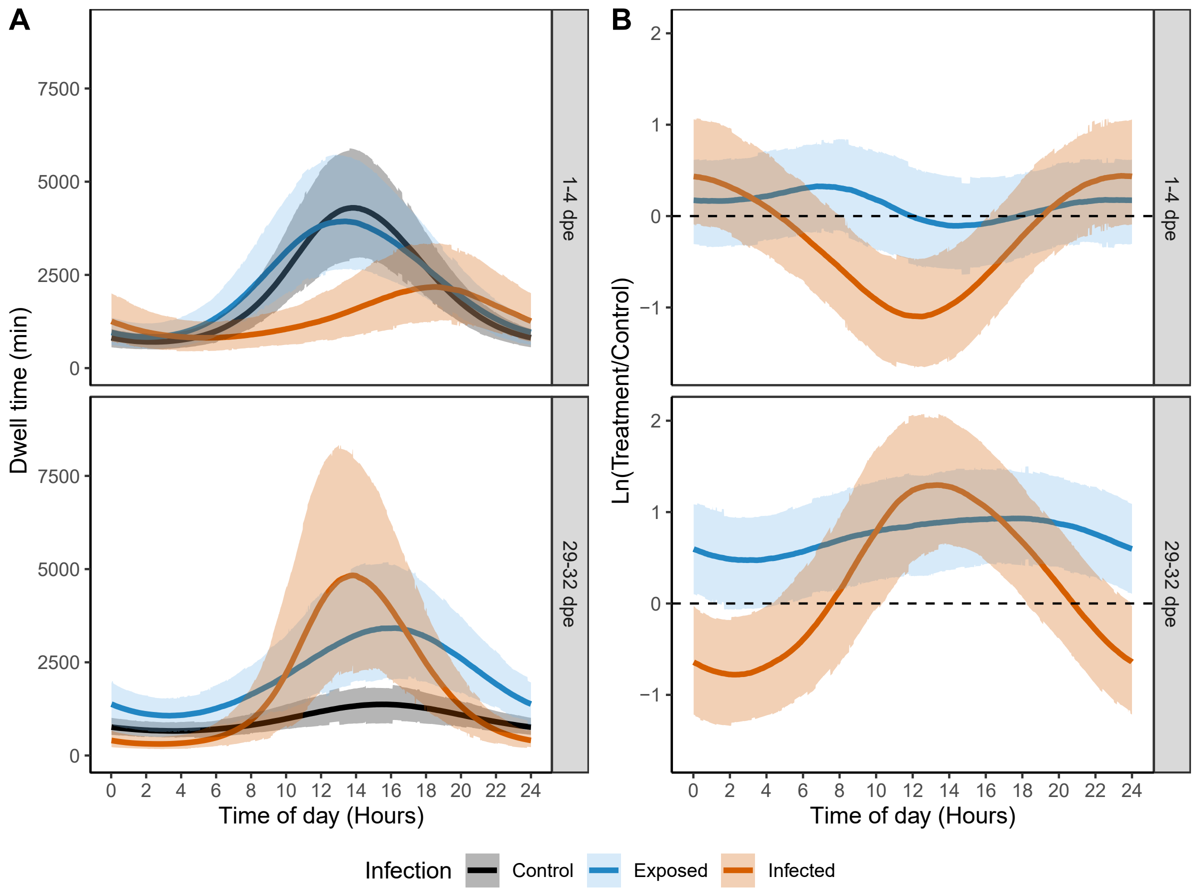


**Supplementary Figure 5. (A)** Expected dwell times (i.e., time spent continuously in one state) as a function of the time of day for *Control*, *Exposed*, and *Infected* fish in state 2 (moderate activity) per recording time. Middle lines display the mean dwell times and upper and lower lines the respective 95% confidence intervals. **(B)** Logarithmic ratio of the deviation in state 2 of exposed and infected fish from the respective control (dashed line) derived from simulations based on the HMM. Middle lines display the means and upper and lower areas the 95% confidence intervals of the distribution.

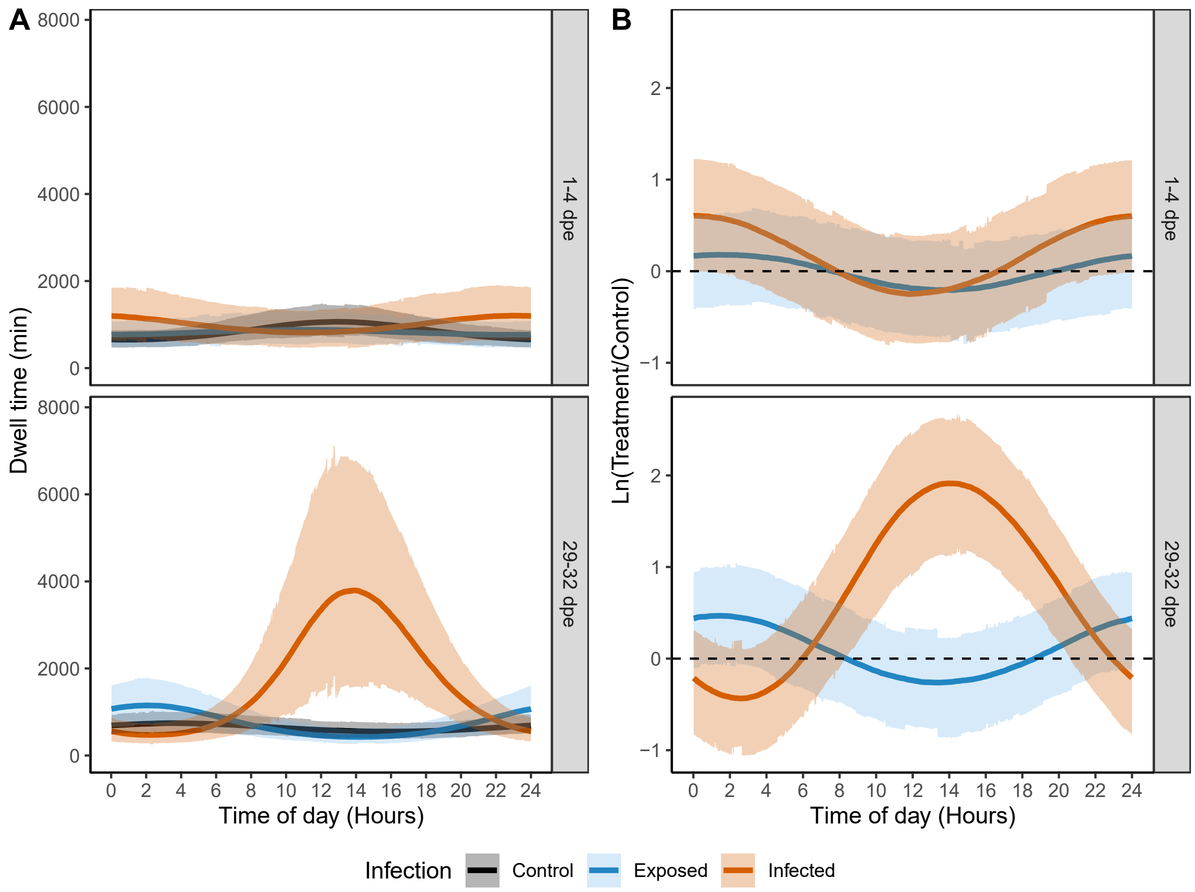


**Supplementary Figure 6. (A)** Expected dwell times (i.e., time spent continuously in one state) as a function of the time of day for *Control*, *Exposed*, and *Infected* fish in state 3 (high activity) per recording time. Middle lines display the mean dwell times and upper and lower lines the respective 95% confidence intervals. **(B)** Logarithmic ratio of the deviation in state 3 of exposed and infected fish from the respective control (dashed line) derived from simulations based on the HMM. Middle lines display the means and upper and lower areas the 95% confidence intervals of the distribution.

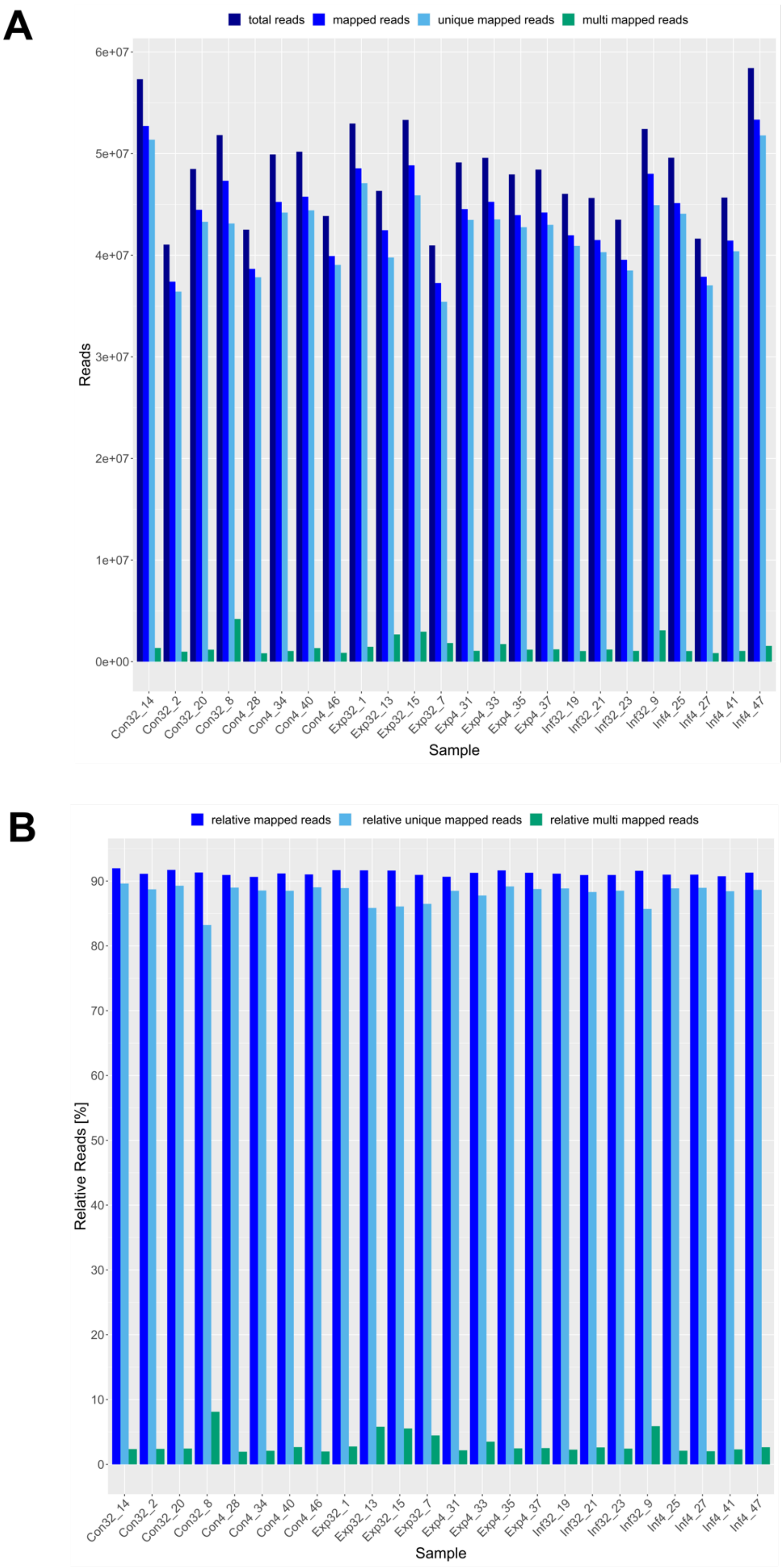


**Supplementary Figure 3: (A)** Total and **(B)** relative number of mapped, uniquely mapped, and multi-mapped reads for all brain samples used for RNA-sequencing. Brain samples originate from *Control* (Con), *Exposed*

(Exp), and *Infected* (Inf) individuals 4 and 32 dpe. The number after the underline represents the individual fish ID
